# Nanoscale imaging resolves canonical topology and intracellular dynamics of SUN5/SPAG4L during mammalian spermiogenesis

**DOI:** 10.64898/2026.03.04.709580

**Authors:** Lukas Herold, Hanna Thoma, Nadja Thielemann, Christian Strissel, Angelina Daube, Silke Braune, Manfred Alsheimer

## Abstract

SUN5 is a testis-specific SUN domain protein essential for connecting the sperm tail to the nucleus. However, until now, its precise localization, intracellular dynamics, and membrane topology during spermiogenesis have remained controversial. To address these discrepancies, we applied ultrastructure expansion microscopy (U-ExM) to systematically track SUN5 redistribution throughout spermiogenesis. This approach enabled a detailed reconstruction of SUN5 localization across developmental stages and revealed previously undescribed enrichment at the perinuclear ring (PNR) and the microtubule manchette, suggesting secondary functions at the PNR or a potential role in intra-manchette transport (IMT). Complementary immunogold labelling using the Tokuyasu method, together with biochemical assays, demonstrated that SUN5 adopts a membrane localization and topology consistent with that of classical SUN domain proteins. Quantitative measurements of the nuclear envelope architecture at the head-to-tail coupling apparatus (HTCA) further enabled us to present a refined structural model of SUN5 positioning at the head–tail junction. Overall, our findings resolve previous discrepancies in the field and provide a coherent framework for understanding SUN5 organization and its role in mammalian spermiogenesis.

**Summary Statement:** In the presented study, we analyzed the dynamic redistribution of SUN5 during mammalian spermiogenesis and resolved its topology in developing spermatids to gain insights concerning the proteins’ molecular function in head-tail coupling.

## Introduction

Nuclear remodeling is a defining feature of eukaryotic cell differentiation, involving extensive chromatin reorganization along with active repositioning and reshaping of the nucleus, both of which are necessary to meet the requirements of the cell in its specific biological context (Burke and Roux, 2009; Skinner and Johnson, 2017). This process is particularly evident during spermiogenesis, a prominent and highly orchestrated phase of male germ cell development in which round spermatids undergo directed morphological transformation into highly specialized spermatozoa. These cells are characterized by specialized organelles, such as the sperm tail and the acrosome, and by a species-specific sperm head distinguished by pronounced nuclear compaction, shaping and elongation (Hermo et al., 2010a; Hermo et al., 2010b; O’Donnell, 2014). Proper nuclear reshaping during sperm differentiation depends on a precisely regulated remodeling of the nuclear envelope (NE) and its essential components (Paci et al., 2018; Pereira et al., 2019).

Germ cell development is tightly coordinated in both temporal progression and structural organization, to ensure the efficient formation of functional sperm (Paci et al., 2018; Manfrevola et al., 2021). A complex structure, known as the head-to-tail coupling apparatus (HTCA), is located at the posterior end of the sperm head and mediates a tight physical connection between the nucleus and the flagellum (Wu et al., 2020). A further key unit is the microtubule manchette, which forms around the posterior part of the nucleus in developing spermatids and mediates sperm head elongation and intra-manchette transport (IMT) of posterior building blocks (Russell et al., 1991; Kierszenbaum, 2002; Kierszenbaum and Tres, 2004; Lehti and Sironen, 2016; Miyata et al., 2024; Gao et al., 2025). It is physically connected to the NE via LINC complexes (Linkers of Nucleoskeleton and Cytoskeleton), evolutionarily conserved protein assemblies that bridge both nuclear membranes (Crisp et al., 2006; Pasch et al., 2015; Gao et al., 2020; Thoma et al., 2023).

LINC complexes are found in nearly all eukaryotic organisms and perform a wide range of functions, most of which are related to their role as platforms for mechanotransduction. They are involved in various processes such as nuclear positioning, anchoring, and reshaping, and they participate in dynamic events including nuclear migration, meiotic telomere attachment and movement, and are central hubs in signaling pathways (Kracklauer et al., 2013; Rothballer and Kutay, 2013; Kim et al., 2015; Lee and Burke, 2018; King, 2023). Structurally, LINC complexes are hexameric assemblies composed of trimeric SUN-domain proteins (Sad1/UNC-84 homology) that interact with trimeric KASH-domain (Klarsicht/ANC-1/Syne1 homology) partners, forming a canonical bridge across the NE. The SUN-domain proteins are located in the inner nuclear membrane (INM), with their N-terminal domains (NTDs) facing the nucleoplasm, while their conserved C-terminal SUN domains extend into the perinuclear space (PNS) via a coiled-coil region. Within the PNS, the SUN domain binds the KASH domain of its partner protein, which is anchored in the outer nuclear membrane (ONM) and exposes its large NTD to the cytosol (Sosa et al., 2012; Wang et al., 2012).

In mammals, five SUN domain proteins have been identified: SUN1 and SUN2, which show widespread expression across most tissues and organs (Hodzic et al., 2004; Padmakumar et al., 2005), and the three SUN domain proteins SUN3, SUN4 (SPAG4), and SUN5 (SPAG4L), which are confined to the testes (Göb et al., 2010; Frohnert et al., 2011; Calvi et al., 2015; Pasch et al., 2015; Shang et al., 2017). SUN3 and SUN4 have been identified as key mediators linking the microtubule manchette to the nucleus. In the context of sperm differentiation, both proteins show nearly identical localization patterns and behavior. Together, they assemble into homo- and/or heterotrimeric complexes that interact with Nesprin1, a connection that appears to be crucial for fertility (Göb et al., 2010; Calvi et al., 2015; Pasch et al., 2015; Gao et al., 2020; Thoma et al., 2023).

While the roles of SUN3 and SUN4 in sperm development are fairly well characterized, the function of SUN5 has remained largely unclear until now. Over the years, independent studies have reported quite conflicting results regarding its localization, its function, and the molecular mechanisms involved. An initial study analyzing SUN5 expression and localization detected the protein at the apical NE facing the acrosome, hinting to a potential role in acrosomal organization or stabilization (Frohnert et al., 2011). An alternative role in meiosis has also been proposed (Jiang et al., 2011; Li et al., 2019), however, Frohnert and colleagues could demonstrate that SUN5 transcripts are absent in the testis until day 15 post-partum, consistent with a strictly postmeiotic expression profile (Frohnert et al., 2011). Later studies were unable to confirm the earlier reports of SUN5 localization to the acrosomal side of the NE. In contrast, Yassine et al. (2015) described a dynamic redistribution of SUN5 during spermiogenesis, detecting the protein in several subcellular compartments before it ultimately accumulated at the head–tail junction (Yassine et al., 2015). A similar distribution pattern was reported shortly thereafter: although in early stages of spermiogenesis SUN5 showed a widespread distribution throughout the cells, followed by progressive enrichment at the HTCA toward the end of the differentiation process (Shang et al., 2017). This latter observation aligns well with several functional studies linking SUN5 mutations to defective head-to-tail coupling in both animal models and human patients (Zhu et al., 2016; Elkhatib et al., 2017; Fang et al., 2018; Alinia et al., 2025). Despite several efforts to characterize SUN5, its reported localization remains inconsistent across studies, and a systematic super-resolution analysis to clarify these discrepancies is still lacking.

While the contribution of SUN5 to HTCA integrity is widely accepted within the field, the exact mechanism of SUN5 function at this site is a matter of debate. In recent years, several models have been proposed to explain the molecular mechanism of SUN5 function, alongside the identification of potential binding partners that might mediate its interactions at the head-tail junction (Frohnert et al., 2011; Zhang et al., 2021b; Zhang et al., 2024; Buglak et al., 2025). Interestingly, some of these models suggest that SUN5 deviates from the canonical topology of SUN domain proteins, for instance by integrating into the ONM with the SUN domain facing the cytosol (Zhang et al., 2021b) or by spanning both nuclear membranes (Zhang et al., 2024). A mechanism involving nuclear pore complexes (NPCs) has also been discussed, as SUN5 has been shown to interact with the NPC component Nup93 (He et al., 2024), and physical connections between NPCs and the cytoskeleton have been reported in *Drosophila* although not involving SUN-domain proteins (Li et al., 2023). Finally, the classical LINC complex model also represents a plausible hypothesis, in which the SUN-domain protein integrated into the INM connects, via an ONM-localized KASH partner, to HTCA associated structures (Sosa et al., 2012; Kracklauer et al., 2013; Rothballer and Kutay, 2013; Kim et al., 2015).

In the present study, we address two central questions regarding SUN5 and its functional role during spermiogenesis. As the currently available data provide limited and partly inconsistent insights into SUN5 distribution, we performed a comprehensive re-evaluation at super-resolution level using ultrastructure expansion microscopy (U-ExM) with a set of newly generated antibodies. Our data strongly refine previous observations and reveal new, formerly undocumented signals. The insights obtained suggest that SUN5 may possess additional functions beyond head–tail coupling and could either contribute to, or be affected by IMT. Next, we determined the precise localization and topology of SUN5 within the NE using our newly generated antibodies against distinct epitopes of the target protein. By applying the Tokuyasu method for immunogold labeling, we generated an extensive set of electron micrographs focusing on the HTCA of elongating spermatids. Quantitative analysis of these micrographs provided strong evidence that SUN5 adopts an orientation within the NE that resembles the canonical topology of SUN domain proteins. Together, our findings shed light on the molecular mechanisms underlying head–tail coupling and open new avenues for future investigations on SUN5.

## Materials and Methods

### Ethics statements

All animal care and experimental protocols were performed according to the guidelines specified within the German Animal Welfare Act (German Ministry of Agriculture, Health and Economic Cooperation). Animal housing and breeding were approved by the local regulatory agency of the city of Würzburg (reference ABD/OA/Tr; according to 111/1 No. 1 of the German Animal Welfare Act). All aspects of mouse work were carried out under strict guidelines to ensure consistent and ethical handling of the mice.

### Animal and tissue preparation

Testis samples and testicular cell suspensions were obtained from 8–20-week-old male mice (*Mus musculus*) of the wild-type C57BL/6J strain or *Sun4* knock-out mutants (*Spag4tm1(KOMP)Mbp*) (Pasch et al., 2015). Animals were euthanized by CO₂ asphyxiation followed by cervical dislocation. Testes were excised and transferred to a glass Petri dish containing ice-cold PBS (140 mM NaCl, 2.6 mM KCl, 6.4 mM Na₂HPO₄, 1.4 mM KH₂PO₄, pH 7.4). After removing the tunica albuginea seminiferous tubules were cut into small pieces and carefully resuspended with a 1ml pipette. The resulting suspension was filtered (30 µm pore size), centrifuged (600 × *g*, 10 min, 4 °C), and resuspended in fresh PBS. For Tokuyasu EM sample preparation, intact tubules were used directly without prior mechanical disruption.

### Antibodies

To generate specific anti-SUN5 antibodies, His-tagged *Sun5* peptides corresponding to the PNS region (AA 95–170, cloned into a pET21a vector) and the NTD (AA 1–61, cloned into a pQE32 vector) of SUN5/SPAG4L2 (genbank reference: NP_001291977) were expressed in E. coli. For the N-terminal construct, the peptide was expressed in tandem, separated by a short linker sequence. The recombinant peptides were purified from the bacteria under denaturing conditions using Ni-NTA agarose columns (Qiagen, Hilden, Germany) and used to immunize guinea pigs or rabbits (SEQLAB Sequence Laboratories Göttingen GmbH, Göttingen, Germany). Antibodies were affinity-purified from the animal sera with the corresponding antigens coupled to HiTrap™ NHS-activated HP columns (Cytiva, Uppsala, Sweden), following the manufacturer’s instructions. Primary and secondary antibodies used in this study are listed in the supplementary information (Table S1, Table S2).

### Ultrastructure Expansion Microscopy (U-ExM)

The U-ExM protocol used in this study was based on the original procedure published by Gambarotto and colleagues (Gambarotto et al., 2019; Gambarotto et al., 2021) and was carefully adapted to meet the specific requirements of the present study.

All incubations were carried out at room temperature (RT) under gentle agitation unless stated otherwise. Poly-L-lysine–coated coverslips (12 mm) were incubated with 500 µl of testicular cell suspension for 20 min on ice. Subsequently, an equal volume of double-strength FA/AA mix (1.4% formaldehyde and 2% acrylamide in PBS) was added. After incubation at 37 °C for 4 h, 40 µl drops of pre-cooled monomer solution (23% sodium acrylate [w/v], 10% acrylamide [w/v], 0.1% N,N′-methylenebisacrylamide [w/v] in PBS) were supplemented with 5 µl of 10% APS and 10% TEMED each and immediately pipetted onto the coverslips. A second coverslip was then placed on top, and gels were allowed to polymerize at 37 °C for 1 h.

Gels were incubated for 15 min in denaturation bufer (200 mM SDS, 200 mM NaCl, 50 mM Tris base, pH 9), detached from the coverslips, and transferred into fresh denaturation buffer. Denaturation was performed at 90 °C for 30 min. Gels were then expanded in ddH₂O, which was replaced twice after 30 min each, followed by overnight incubation at RT.

The following day, gels were washed three times in PBS for 15 min each and subsequently incubated in 0.1% Triton X-100 in PBS for 20 min. Blocking was performed for 1 h using a dedicated blocking buffer (5% milk, 5% FCS, and 0.1% Tween 20 in PBS). Primary antibody incubation was carried out overnight at 4 °C in PBT, followed by three washing steps—two in PBST (0.1% Tween 20 in PBS) for 15 min each and one in PBS for 15 min. Secondary antibodies were incubated for 3 h at RT and washed as described above. Finally, gels were expanded in ddH₂O three times for 15 min each.

Expanded gels were mounted on silanized coverslips and placed into custom imaging chambers, remaining submerged in ddH₂O during imaging. Imaging was performed using either a Leica SP2 confocal laser-scanning microscope (Leica Microsystems GmbH, Wetzlar, Germany) equipped with a 63× oil-immersion objective or a Leica Thunder widefield imaging system (Leica Microsystems GmbH, Wetzlar, Germany) with a 100× oil-immersion objective for 3D analysis. Images were deconvolved with Huygens Essential 25.04 software (Scientific Volume Imaging, Hilversum, Netherlands) using standard settings for confocal microscopy and theoretical points-spread function (PSF).

### Transmission electron microscopy (TEM) according to Tokuyasu

Tokuyasu electron microscopy was performed following previously established protocols (Slot and Geuze, 2007; Möbius and Posthuma, 2019; Link et al., 2024)

All incubations were carried out at room temperature (RT) under gentle agitation unless stated otherwise. In brief, freshly extracted testicular tubules were transferred to reaction tubes and fixed with 2% formaldehyde (FA) in 0.1 M PHEM buffer (60 mM piperazine-N,N′-bis(2-ethanesulfonic acid) [PIPES], 25 mM 4-(2-hydroxyethyl)-1-piperazineethanesulfonic acid [HEPES], 10 mM ethylene glycol-bis(β-aminoethyl ether)-N,N,N′,N′-tetraacetic acid [EGTA], 2 mM MgCl₂, pH 6.9 adjusted with 1 M NaOH) for 10 min. The fixative was replaced once, and incubation was continued for an additional 2 h.

Samples were rinsed three times with 0.1 M PHEM, quenched with 0.15% glycine in 0.1 M PHEM for 10 min, and gently pelleted by centrifugation. Next, 200 µl of 12% gelatin in 0.1 M PHEM was added, and the tubes were placed in a pre-heated centrifuge equipped with a swing-out rotor and spun at 2500 × *g* for 2 min at 37 °C. The supernatant was removed, replaced with fresh gelatin solution, and spun again before allowing the block to harden on ice for 30 min. The tubes were then opened at 4 °C, and the pellet was cut into small blocks (∼2 mm diameter), which were subsequently immersed in 2.3 M sucrose in 0.1 M PHEM and incubated at 4 °C overnight. Blocks were mounted on aluminum pins using 2.3 M sucrose as adhesive and rapidly frozen in liquid nitrogen.

Cryo-sectioning was performed with a Leica EM UC7 ultramicrotome (Leica Microsystems GmbH, Wetzlar, Germany) using diamond knives optimized for cryo-sectioning (DiATOME Ltd., Nidau, Switzerland). Ultrathin sections (80 nm) were cut at -120 °C, collected using a wire loop with pick-up solution (2.3 M sucrose and 1.8% methyl cellulose in 0.1 M PHEM), transferred onto nickel grids, and thawed.

For immunogold labeling, the pick-up solution was removed by washing the sections in PBS for 30 min. Sections were then incubated twice with 0.15% glycine in PBS for 5 min each and blocked with 1% BSA in PBS for 10 min. Primary antibodies were diluted in antibody dilution buffer (0.1% BSA and 0.5% fish gelatin in PBS) and incubated for 1.5 h at RT. Samples were washed five times in 1% BSA in PBS for 3 min each. Secondary antibody incubation was performed under identical conditions. Sections were then washed three times with 1% BSA in PBS and twice with PBS before being post-fixed in 1% glutaraldehyde (GA) in PBS for 5 min. After rinsing three times in ddH₂O for 5 min each, grids were stained with 2% uranyl acetate in ddH₂O for 10 min and then briefly rinsed in ddH₂O before final contrasting in a 1:1 mixture of 1.8% methyl cellulose and 0.4% uranyl acetate for 10 min on ice. Excess contrasting solution was removed by gently dragging the grids over filter paper within a pick-up loop, and the grids were allowed to dry for at least 30 min before imaging.

Electron micrographs were acquired using a JEOL JEM-1400 Flash scanning transmission electron microscope (STEM) (JEOL Ltd., Akishima, Japan) operated at 120 kV acceleration voltage with a Matataki Flash 2k × 2k camera system. Alternatively, a JEOL JEM-2100 transmission electron microscope (TEM) (JEOL Ltd., Akishima, Japan) equipped with a TVIPS F416 4k × 4k camera system (Tietz Video and Image Processing Systems GmbH, Gilching, Germany) was used to record high-magnification images at 200 kV.

### Image analysis

Imaging data were analyzed using Fiji as imaging-processing platform (version 1.54f, (Schindelin et al., 2012)). For specific analytical tasks (see below) custom macros were designed by the authors and implemented with the assistance of the AI tool ChatGPT (GPT-5, OpenAI, San Francisco, CA, USA; accessed October 2025). Designed macros were reviewed, tested, and validated by the authors prior to analysis. Macro scripts and parameter settings are provided in the supplementary information Code S1 and Code S2.

Immunogold localization experiments were evaluated by marking the center of the PNS with the segmented line tool and gold particles were classified into cytoplasmic and nuclear pools based on their position relative to the PNS center. Running the macro automatically measured the distance of each gold particle to the PNS center and compiled the results into a table, assigning negative values to particles present on the nuclear side of the NE. This procedure was performed across multiple images, and only regions in which both nuclear membranes were clearly resolved were included in the analysis.

Measurements of the distance between the two nuclear membranes were obtained from ultra-closeups of the HTCA. Image regions were selected according to the same criteria as above. A linear ROI was drawn perpendicular to the membranes approximately within the relevant region. Running the macro generated a Gaussian-smoothed k-plot corresponding to the intensity along the linear ROI, form which the peak-to-peak distance was automatically measured and recorded. To visually validate each measurement, the resulting plot was displayed to the user along with the measured distances. This process was repeated several times with linear ROIs spaced about 5 nm from each other to create the full dataset.

### Statistical analysis

Statistical analyses were performed using OriginPro 2023 (OriginLab Corp., Northampton, MA, USA). For gold localization experiments, data were combined into two main datasets (NTD and PNS, respectively) containing the measured distances to the PNS center. Normality of the distributions was assessed using the Shapiro–Wilk test, and variances were compared with an F-test. Group differences were evaluated using Student’s t-test, which was safeguarded by Welch’s t-test. Effect size was determined by calculating Cohen’s *d* value. Following statistical evaluation, the data were visualized using boxplots and plots displaying 95% confidence intervals.

### Alpha Fold Structure Predictions

Structural predictions of trimeric SUN5 assemblies were generated using AlphaFold2-multimer (v3), employing as input the sequence region of SUN5/SPAG4L2 spanning from the transmembrane domain to the C-terminus of the protein (AA 103-373). The highest-ranking model (model_2, seed_000) exhibited a mean pLDDT of 80.8 and predicted TM-scores of pTM = 0.744 and ipTM = 0.731, indicating high confidence in both overall fold and subunit interfaces. Model with local pLDDT values can be found in Figure S1.

### Generation of plasmid constructs and transfected culture cells

To generate plasmid constructs coding for Myc-SUN5-FL-EGFP and Myc-SUN5-ΔC-EGFP, SUN5-FL and SUN5-ΔC cDNAs were amplified by RT-PCR from RNA isolated from mouse testis suspensions with respective primers designed based on the long isoform of SUN5 (SPAG4L2; genbank accession number NM_001305048). PCR products were cloned into a modified pCMV-Myc-EGFP vector (Thoma et al., 2023) in frame to the Myc- and EGFP-tags, and sequenced for verification.

COS-7 (DSMZ; ACC-60) were grown in Dulbeccós modified Eaglés medium (DMEM; Gibco by Life Technologies, Darmstadt, Germany), into which 10% fetal calf serum (FCS; Capricorn Scientific, Ebsdorfergrund, Germany) and 1% penicillin-streptomycin (Thermo Fisher Scientific, Dreieich, Germany) were added. Cultures were grown at 37°C and 5% CO_2_ concentration. Cells were transfected with the Myc-EGFP constructs using PolyMag transfection reagent (OZB-IV-TK30210; OZ Biosciences, San Diego, CA, USA) according to the manufacturer’s protocol. Transfection was repeated once after 1 h on the same cells. Cells were subsequently cultured at 37°C (5% CO_2_) and analyzed 24 h after final transfection.

### *In situ* proteinase K digestion assay

*In situ* proteinase K digestion assays were conducted as previously described (Thoma et al., 2023). In brief, NIH-3T3 culture cells were transfected in 35 mm culture dishes with double-tagged SUN5 constructs (N-teminal Myc and C-terminal GFP-Tag according to the assays previously described for SUN4 (Thoma et al., 2023). 24 h after transfection, each well was washed twice with ice-cold KHM-buffer (110 mM KOAc, 2 mM MgCl_2_, 20 mM HEPES, pH 7.4) for 1 min each. For each construct, one well was permeabilized with 24 µM ice-cold digitonin on ice for 10 min, followed by two washes with ice-cold KHM bufer (1 min each) to remove the reagent. Subsequently, proteinase K was added at a final concentration of 4 µg/ml. A second well received 4 µg/ml proteinase K without prior permeabilization, while a third well was treated with 4 µg/ml proteinase K in the presence of 0.5% Triton X-100. A fourth well served as an untreated control and was incubated with KHM bufer only. The cells were incubated 45 min at RT before the reaction was stopped by adding proteinase inhibitor cocktail (Roche Diagnostics GmbH, Manheim, Germany). Cells were harvested using a cell scraper, transferred into reaction tubes and washed in KHM.

Finally, cells were resuspended in SDS-sample buffer (120 mM Tris-HCl pH 6.8, 10% SDS, 20% glycerol, 20% 2-mercaptoethanol and Bromophenol Blue) and incubated at 95°C for 5–10 min. Samples were subjected to SDS-PAGE followed by Wester blotting onto nitro-cellulose membranes and were probed with anti-Myc and anti-GFP antibodies, respectively.

### Manuscript preparation

We used the AI tool ChatGPT (GPT-5, OpenAI, San Francisco, CA, USA; accessed January 2026) for minor language refinements and stylistic editing during manuscript preparation.

## Results

For detailed analysis of SUN5 localization and dynamics during spermiogenesis, we conducted several antibody dependent experiments including U-ExM across multiple cellular stages, Tokuyasu EM gold immunolocalizations and digitonin based topology assays. To this end, we generated custom antibodies against the N-terminal domain and a region of the C-terminal domain (PNS region) excluding the SUN domain, respectively. For quality check of these antibodies, we performed Western Blot experiments on testes tissue lysates (Fig. 1B) and double-localization U-ExM experiments with both the N-terminal and the PNS directed SUN5 antibodies, respectively, as well as separate co-stainings with anti-SUN4 antibodies as a reference (Fig 1C-E).

**Figure 1:**
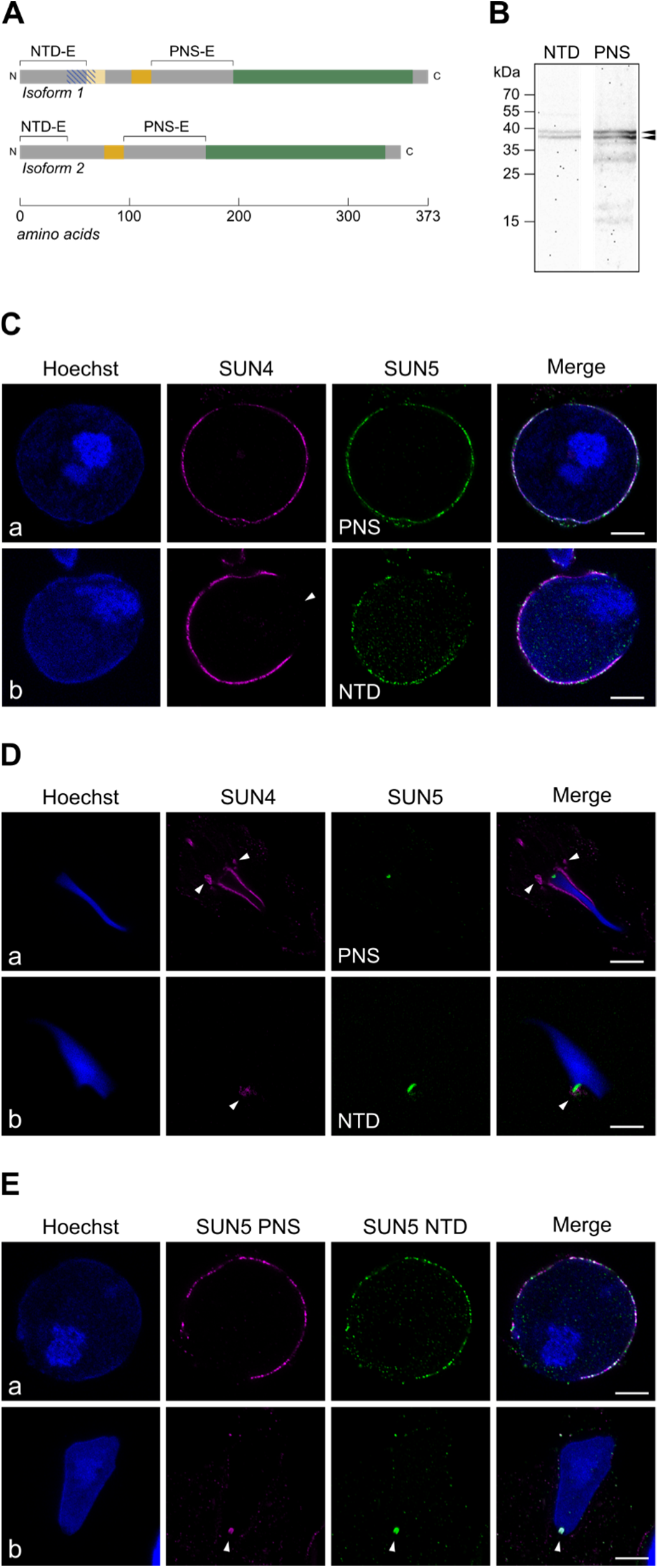
Immunofluorescence analysis with custom antibodies against two distinct SUN5 epitopes. (A) Schematic representation of the two murine SUN5 isoforms (isoform1: SPAG4L-2; isoform2: SPAG4L) and the regions used for antibody generation. The N-terminal domain (NTD) epitope (NTD-E), the putative perinuclear space (PNS) epitope (PNS-E), the SUN domain (green), the TM domain (orange), and hydrophobic domain of isoform 1 (light orange) are indicated. The blue hatched area marks the isoform 1 specific amino acids. (B) Western Blot of whole testis lysates probed with anti-SUN5-NTD and anti-SUN5-PNS antibodies, respectively. Specific signals at approximately 37 kDa corresponding to the two SUN5 isoforms, Spag4L and Spag4L2, are indicted with arrowheads. (C) U-ExM immunolocalizations with either of the two SUN5 antibodies on early round spermatids counterstained with anti-SUN4 antibodies. Both the anti-SUN5-PNS (a) and the anti-SUN5-NTD (b) exhibit similar distribution patterns, including their exclusion from the acrosomal side of the nucleus (white arrowhead in b). (D) Corresponding U-ExM immunolocalizations on late elongating spermatids. While SUN4 accumulates to the redundant nuclear envelope (white arrowheads), the SUN5 signal detected by both antibodies accumulates at the head–tail junction. (E) Double immunolocalizations using both SUN5 antibodies show a high degree of overlap in (a) early round spermatids and (b) late elongating spermatids. Scale bars: 10 µm; Images are representative of at least four experimental repeats

As expected for antibodies directed against the same protein, the anti SUN5-NTD and anti SUN5-PNS antibodies exhibited highly concordant signal patterns in both early round and late elongated spermatids. Co-staining experiments confirmed this observation by demonstrating a substantial overlap of the two signals (Figure 1E). Both antibodies directed against SUN5 labelled the nuclear envelope in early round spermatids and a distinct region of the implantation fossa in late elongated spermatids. Localization of SUN5 to the nuclear envelope in round spermatids was further corroborated by counterstaining with SUN4, a well-established marker of the INM (Pasch et al., 2015; Thoma et al., 2023). In late elongated spermatids, however, SUN5 was exclusively observed at the head–tail junction in clear contrast to the distribution of SUN4, which accumulates in the redundant nuclear envelope of elongated spermatids and is subsequently removed from the final spermatozoa (Fig 1 D, E). Together, these results validated the suitability of the newly generated antibodies for use in subsequent experiments.

### SUN5 adopts a canonical topology of SUN domain proteins

As summarized above, previous studies have reported conflicting results regarding the membrane topology of SUN5, with key aspects of their conclusions being mutually exclusive. Most recently, Buglak et al. (2025) proposed three different models for the positioning of SUN5 within the nuclear membrane. However, it remains unclear which model most accurately reflects the true localization and membrane topology of SUN5. To address this, we performed detailed electron microscopical analysis on testis tissue sections by applying the Tokuyasu method (Tokuyasu, 1973; Möbius and Posthuma, 2019) for immunogold labeling. The Tokuyasu method has, to our knowledge, not yet been applied to structures of the HTCA region in mammalian spermatids and, therefore, we initially pre-evaluated the ultrastructural preservation of this specific region in samples prepared according to this protocol to ensure that the observed localizations accurately reflect the *in vivo* situation.

As can be clearly seen in Fig. 2A, the electron microscopic images show very good structural preservation of the HTCA region and the adjacent nuclear membranes in elongating spermatids. All key components revealed their typical shape, size, and relative positions to each other, including the proximal (PC) and distal (DC) centrioles, the segmented column complex (SCC), the capitulum (Ca), and the basal plate (BP) (for comparison see Wu et al., 2020). The SCC structure appeared highly pronounced and showed the characteristic periodic striation, indicating adequate contrasting and structural integrity. Most importantly and essential for the downstream analysis, both the inner and the outer nuclear membranes were also well resolved, visible as white lines that are separated by the PNS, which appeared as a darker-stained area between the INM and ONM (Fig. 2A, B). Together, these observations demonstrated that the ultrastructure was well preserved, allowing the following immunogold localization experiments to be considered reliable.

**Figure 2:**
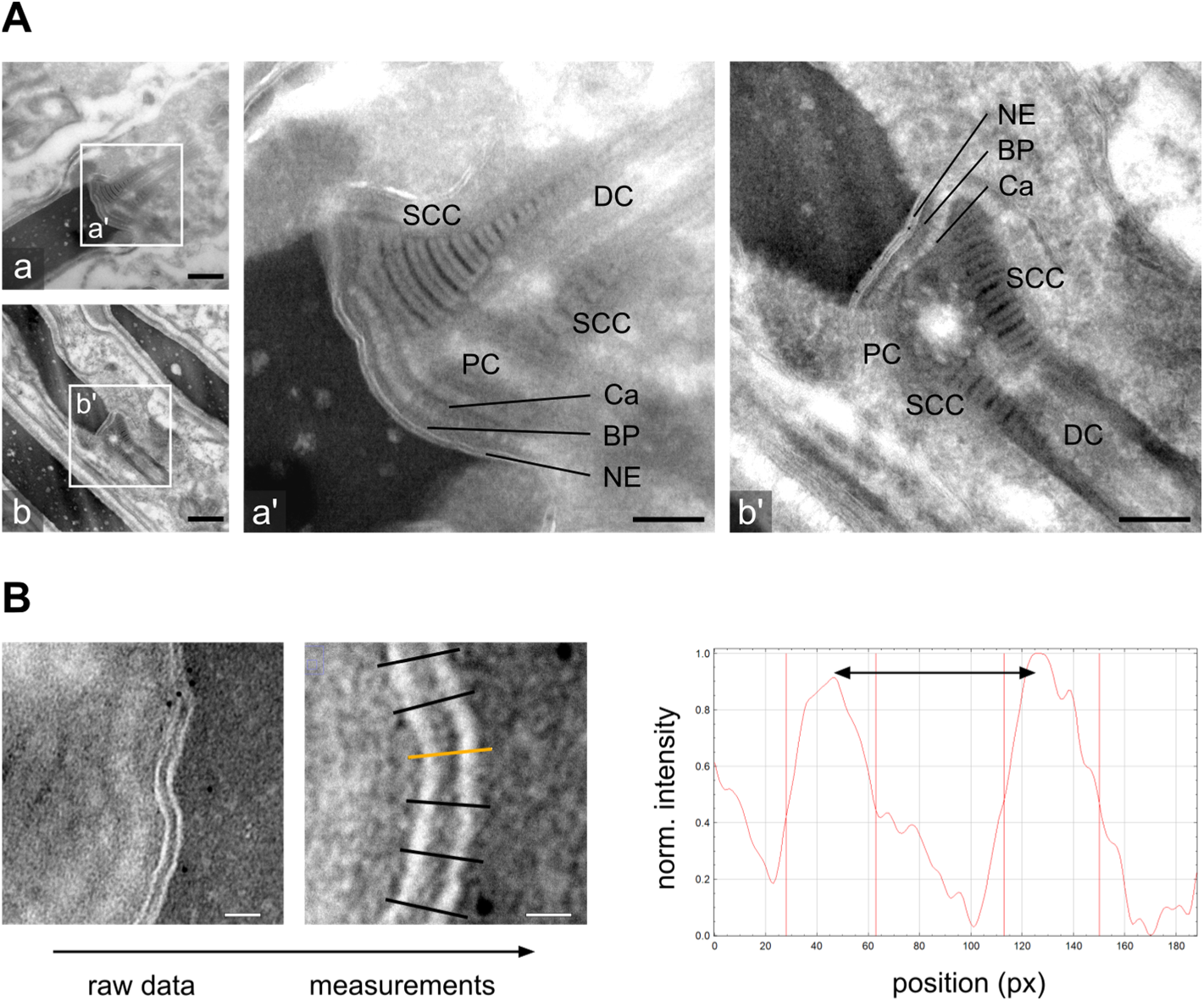
Ultrastructural evaluation of Tokuyasu EM samples and measurement of the INM-ONM distance. (A) Overview images (a, b) of elongating spermatids (scale bars: 500 nm). Higher-magnification images (a’, b’) of the highlighted regions in a and b with annotated ultrastructural features (scale bars: 200 nm). Depicted are the segmented column complex (SCC), the distal (DC) and the proximal centriole (PC), the capitulum (Ca), the basal plate (BP) and the nuclear envelope (NE). (B) Workflow for measuring the INM-ONM distance (scale bar: 20 nm). Images are representative of at least two experimental repeats.

Before collecting quantitative data concerning SUN5 orientation and localization, we measured the distance between the nuclear membranes, allowing us to generate a realistic model of SUN5 during the subsequent analysis. To this end, we took ultra–close-up images of the NE at 80 k magnification. The resulting imaging data was then quantified using a Fiji macro developed for this purpose (Code S1), which used a gaussian smoothed peak detection in intensity profiles along linear ROIs to pin down the distance. This analysis revealed that the membranes were on average separated by 14.25 nm (n = 100, SD_NE_ = 3.16 nm, SE_NE_ = 0.32 nm).

Using the two antibodies directed against the N-terminal domain and an epitope between the TM and the SUN-domain, respectively, we then set out to define the protein’s topology within the NE at the HTCA region. Immunogold staining was performed to determine whether SUN5 resides within the INM or the ONM and to clarify its actual orientation in the membranes. For both antibodies a pronounced accumulation of gold particles was observed at the nuclear envelope in the HTCA region, with only minimal background signal in the surrounding areas (Fig. 3A). With antibodies against the N-terminal domain (see Fig. 3A a-b) we observed gold particle enrichment at the INM, with a clear preference for the nuclear side of the membrane. Gold particles directed against the putative PNS-associated epitope were found dispersed across the PNS, showing a tendency toward the center between the inner and the outer nuclear membranes (Fig 3A c-d). In both instances, some particles deviated from the expected localization pattern; however, such variation is anticipated when working with sandwich antibody systems at this spatial scale.

**Figure 3:**
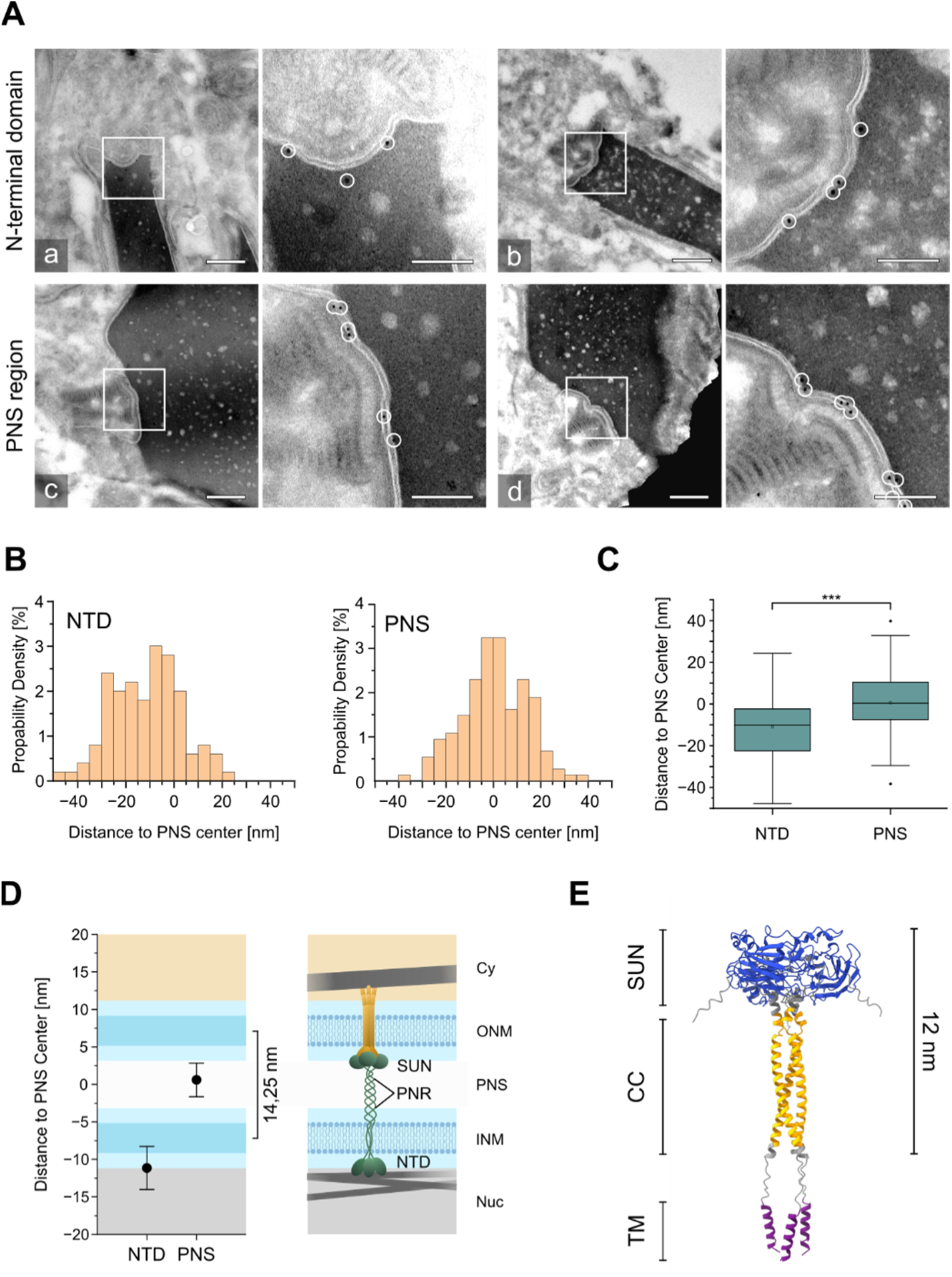
Immunogold-based topology assay and model of SUN5 within the nuclear envelope. (A) Representative images of anti-SUN5-NTD (a, b) and anti-SUN5-PNS (c, d) immunogold labeling at the HTCA of elongating spermatids. Overview images are shown on the left (scale bars 500 nm) with corresponding higher magnifications on the right (scale bars 200 nm). Gold particles are encircled for enhanced visibility. (B) Histograms of the analyzed gold particles (anti-SUN5-NTD: n = 100; anti-SUN5-PNS: n = 148) relative to their distance to the PNS center. (C) Box plot summarizing the immunogold dataset. Statistical significance was assessed using an Student’s t-test (p < 0.0001) (D) Left: 95 % confidence intervals of the immunogold dataset plotted onto a plotting area representing the NE (left). Highlighted are the cytosol (Cy) in orange, the two nuclear membranes (INM and ONM), separated by a measured distance of 14,25 nm, in blue with the darker shade representing the lipid bilayer. The nucleoplasm (Nuc) is depicted in grey. Right: Schematic model of SUN5 as trimeric assembly (green) within the NE; a hypothetical trimeric binding partner residing in the ONM is shown in orange. (E) Alpha-fold based structural prediction of a SUN5 trimeric assembly form the transmembrane domain to the C-terminal end. The SUN domain is shown in blue, the coiled-coil region (CC) in yellow and the trans membrane domain (TM) in purple.

To pinpoint the precise location of the SUN5 N-terminus and its putative PNS region relative to the nuclear membranes, we performed multiple measurements across several cells (n > 50). Using a self-generated Fiji macro (Code S2) we quantified the distribution of gold particles based on their displacement from the PNS center, with negative values indicating a displacement from the PNS center toward the nucleus and positive values representing displacement toward the cytosol. Statistical analysis revealed mean values of –11.15 nm (n = 100 particles) in case of the SUN5-NTD, strongly suggesting that the NTD is orientated towards the nucleus. Due to the linkage error, standard deviations of these measurements where expectedly high (SD_SUN5-NTD_ = 14.39 nm) but standard errors were low owing to the large amount of collected data (SE_SUN5-NTD_ = 1.44 nm). On the other hand, a mean value of 0.59 nm (n = 148 particles, SD_SUN5-PNS_ = 13.70 nm, SE_SUN5-PNS_ = 1.13 nm) was calculated for the SUN5-PNS samples, indicating that the presumed PNS epitope is indeed located within PNS. Statistical analysis, including Students’ t-test confirmed a highly significant (p < 0.0001) difference between the groups, which is also apparent in the resulting box plots (Fig. 3 B-C).

The measured distance between the nuclear membranes and a value of lipid bilayer thickness of 4 nm (Mitra et al., 2004) were then used to define a zoned graphing area in which the 95% confidence intervals of the gold localization data were plotted (Fig. 3D). The resulting graph confirmed the visual inspections and demonstrates that the SUN5 NTD is located on the nucleoplasmic side of the INM, while the PNS region localizes within the PNS, consistent with the canonical topology of SUN-domain proteins. Based on the TEM data, a model depicting SUN5 in the NE was generated to illustrate its topology (Fig. 3D). AlphaFold predictions of trimeric SUN5 assemblies, generated for the region spanning the transmembrane domain to the C-terminal end while excluding the unstructured N-terminus, aligned well with this model. In this prediction, the structured coiled-coil domain together with the conserved SUN domain span only approximately 12 nm (Fig. 3E), suggesting that SUN5 cannot bridge both membranes.

### Biochemical confirmation of canonical SUN5 topology

To verify our findings regarding the topology of SUN5, we performed a biochemical approach based on proteinase K digestion, as previously described (Thoma et al., 2023). For this assay, we expressed Myc- and EGFP-tagged SUN5 fusion proteins in Cos-7 cells, either as full length or C-terminally truncated versions. Using digitonin, we selectively permeabilized the plasma membrane of the cells while leaving organelle and nuclear membranes intact. Subsequent treatment with proteinase K was expected to digest the exposed cytoplasmic and nucleoplasmic domains of the proteins, while domains located inside endogenous membrane systems, like the ER or the perinuclear space, should remain protected (Thoma et al., 2023).

Using this assay, we first tested the double tagged (NTD: Myc-tag, C-terminal domain (CTD): EGFP) full-length SUN5 construct. As shown in Fig. 4, the anti-Myc and anti-GFP antibodies both detected the undigested Myc/EGFP-tagged full-length protein (apparent molecular weight ∼75 kDa) in untreated as well as in proteinase K-treated cells with intact membrane systems. When cells were treated with 24 µM digitonin, the Myc signal almost completely disappeared, indicating that the NTD became accessible to the enzyme upon permeabilization of the plasma membrane. In contrast, the C-terminal EGFP signal remained detectable but exhibited a downshift of approximately 14 kDa, which corresponds to the size of the Myc-tagged NTD. Samples treated with Triton X-100 during the digestion showed no detectable signal for either the Myc-tag for the EGFP-tag, demonstrating complete permeabilization of all cellular membrane systems. Notably, the proteinase K positive control did also exhibit small amount of down-shifted signal, likely reflecting partial cell damage introduced during sample processing.

**Figure 4:**
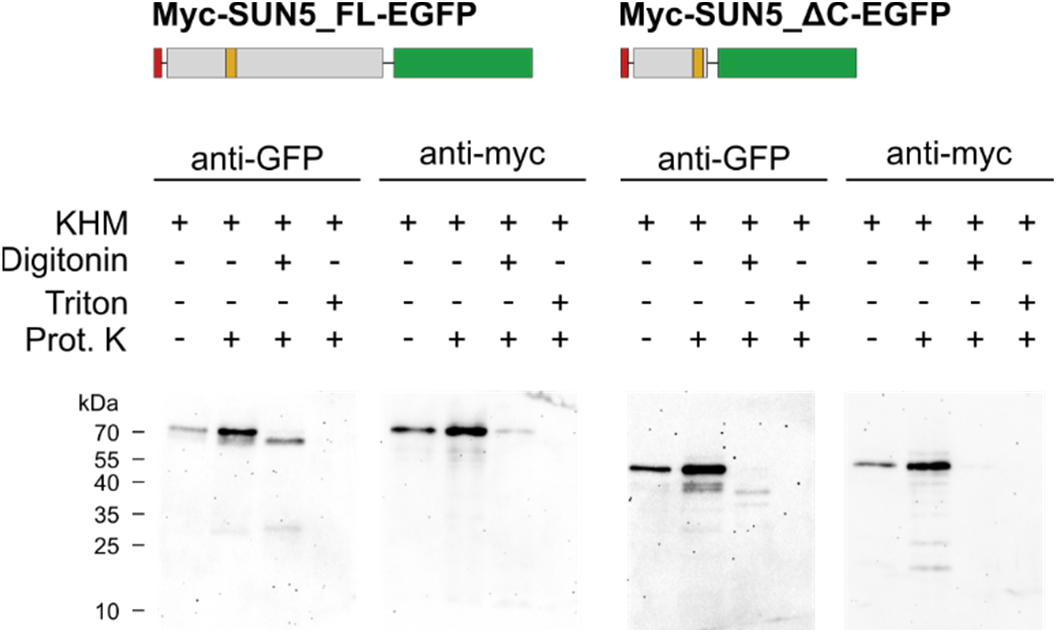
Identification of SUN5 topology by *in situ* proteinase K digestion assay. COS7 cells expressing double tagged SUN5 constructs were treated with either digitonin or Triton X-100 for selective permeabilization and subjected to proteinase K digestion. Along with negative controls, samples were analyzed by SDS-PAGE and subsequent Western blotting. Probing with anti-GFP and anti-Myc antibodies revealed a downshift of the C-terminal EGFP signal upon digitonin permeabilization of the cells, indicating that the C-terminal SUN domain is protected within the PNS, while the N-terminal domain appears to be exposed to the cytoplasm and susceptible to proteolytic digestion. Images are representative of at least two experimental repeats.

Virtually identical results were obtained with the C-terminally truncated SUN5 construct (Fig. 4). Signals in control cells, both untreated and treated with proteinase K, appeared at approximately 46 kDa when probed with anti-Myc and anti-GFP antibodies, consistent with the predicted molecular weight of the Myc- and EGFP-tagged C-terminal deletion construct. As observed for the full-length construct, treatment with 24 µM digitonin followed by proteinase K digestion, resulted in a downshift of approximately 14 kDa to an apparent molecular weight of approximately 32 kDa when probed with the anti-GFP, suggesting a loss of the N-terminal domain by proteolytic digestion. In contrast, no signal could be detected with the anti-Myc-antibodies under the same condition. As expected, the addition of Triton X-100 led to complete degradation of the protein.

Together, these results again support the hypothesis that SUN5 shows a canonical SUN domain protein topology.

### Dynamic redistribution of SUN5 during sperm differentiation

While elucidating the topology at the HTCA is a crucial part for understanding the role of SUN5 in the context of head-tail coupling, we extended our analysis to obtain a more holistic picture of SUN5 during the spermiogenesis process. To this end, we next performed a comprehensive analysis of SUN5 localization and behavior by applying U-ExM not only at selected time points but across the entire span of spermiogenesis, with a particular focus on intermediate stages. We examined SUN5 distribution in super-resolution to pin down new aspects and previously unrecognized features.

Initial observations in early round spermatids revealed weak, but specific signals that appeared outside the nuclei with a tendency to accumulate in one distinct region of the cell. Double staining experiments together with GM130, a marker of the cis-Golgi network (Nakamura et al., 1995) showed that these signals exhibit clear spatial association to the GM130 signal (Fig. 5A). However, exact co-localization was only rarely observed, and in most cases the SUN5 signal appeared slightly displaced from that of GM130, indicating an accumulation of SUN5 in the trans-Golgi network. In addition to the cytoplasmic signal, we found SUN5, similar to the INM component SUN4 (Thoma et al., 2023), also localizing to the NE from this stage onward. In contrast to the initial characterization of SUN5 (Frohnert et al., 2011), we did not detect SUN5 at the acrosomal side of the NE, but instead at the lateral and posterior regions of the nucleus (Fig. 5B,C). Our observations are largely consistent with the report by Yassine et al. (2015). However, in contrast to our findings here, they reported that SUN5 accumulates preferentially in the cis-Golgi rather than in the trans-Golgi network in early round spermatids.

**Figure 5:**
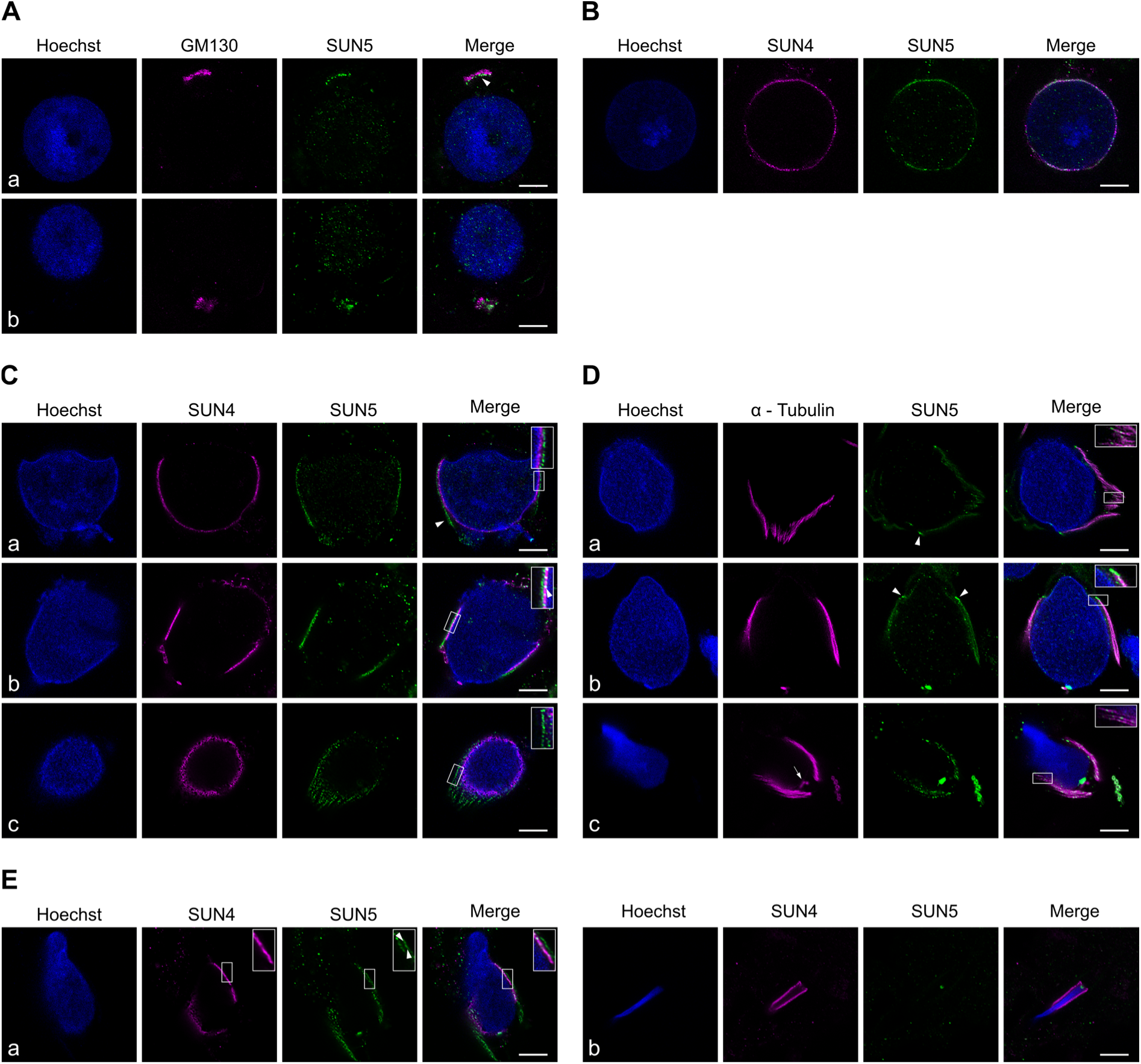
U-ExM analysis of the dynamic redistribution of SUN5 throughout spermiogenesis. (A) SUN5 signal closely resembles the distribution of the cis-Golgi marker GM130. A small spatial ofset is frequently observed (white arrowhead in merge a). (B) In round spermatids, SUN5, similar to SUN4, accumulates at the NE. (C) In progressing spermatids, the localization patterns of SUN5 and SUN4 diverge (enlarged box in a), which is particularly evident toward the posterior cell pole (white arrowhead in a). Two distinct pools of SUN5 are observed (white arrowhead in the enlarged box in b). Dotted SUN5-positive structures extend from the nucleus towards the posterior cytosol without associating with the SUN4 signal (c). (D) In progressing spermatids counterstained for α-tubulin, an association of SUN5 with the microtubule manchette becomes apparent. In addition, a bright signal at the anterior edge indicates association with the PNR (arrowheads in a and b). Enlarged boxes in a and c highlight the tendency of SUN5 to colocalize with α-tubulin. Accumulation of the SUN5 signal near the centriole pair and thus the HTCA is also frequently observed (white arrow in c). (E) SUN5 distribution in later stages of spermiogenesis. In elongating spermatids, SUN5 remains detectable in two distinct pools (white arrowheads and enlarged box in a) as well as at the HTCA. In very late stages of elongation, only the HTCA signal persists, whereas the peripheral signal disappears (B). Scale bars: 10 µm; Images are representatives of at least four experimental repeats. Shown are results produced with both the anti-SUN5-NTD antibodies (A; C; D, c; E, a) and the anti-SUN5-PNS (B; D, a,b; E, b), which showed similar distribution patterns in each of the differentiation steps.

Following early round spermatids, we then evaluated SUN5 signal distribution in more advanced stages, which were previously reported to show SUN5 in the posterior half of the NE before accumulating at the sperm neck region (Yassine et al., 2015; Shang et al., 2017). Consistent with this, our U-ExM analysis revealed SUN5 signal in the NE along the nuclear periphery extending towards the posterior cell pole, which was corroborated by co-staining of SUN4 (Fig. 5C). A gradual accumulation of signal at the HTCA was detectable from stage seven of spermiogenesis onwards and increased significantly in intensity during development (Fig. 5D-E). Interestingly, more detailed analysis of the U-ExM samples revealed that during sperm head elongation, the peripheral nuclear signal resolved into two distinct pools. One pool localized directly adjacent to the chromatin, consistent with a NE localization, while the second pool was positioned spatially separated from the INM marker SUN4 toward the cytoplasm (Fig 5C). This displacement was already detectable in lateral nuclear regions and became more pronounced toward the posterior cell pole. In this region, SUN4 curved along the NE outlining the nuclear staining, while the SUN5 signal separated from the nuclear periphery and extended towards the posterior cell pole in linear projections. These signals are characterized by a punctate appearance and a distribution that closely resembles the shape and extent of the microtubule manchette, which attaches to the PNR and extends toward the posterior pole (Lehti and Sironen, 2016).

To further analyze this phenomenon, U-ExM was performed using anti-SUN5 antibodies in combination with an α-Tubulin staining (Fig. 5D). This analysis revealed a clear association between the SUN5 peripheral signal and the microtubule manchette, observed with both the NTD and the PNS domain directed antibodies. Notably, in elongating spermatids, SUN5 accumulated at the anterior edge of the manchette without direct colocalization to α-Tubulin, suggesting that it specifically localizes to the PNR or associated structures (Fig. 5D; Fig. S2; Movie S1; see below).

At late stages when the elongation process was nearly complete, the SUN5 manchette associated signal disappeared, coinciding with manchette disassembly at step 14 of spermiogenesis (Lehti and Sironen, 2016). Following this event, only the bright, spot-shaped signal persisted at the HTCA region, exhibiting strong intensity (Fig. 5E).

To summarize and illustrate these findings, a schematic model was generated (Fig. 6). According to our data, SUN5 is initially detected in the trans-Golgi network and then gradually accumulates at the nuclear envelope, where it first distributes across the entire nuclear periphery but becomes progressively confined to the posterior cell pole as acrosome development progresses. During step 7-10 of spermiogenesis the SUN5 pool within the NE overlapping with that of SUN4, becomes progressively reduced. Concurrently, a second pool of SUN5 emerges that is spatially distinct from the NE, but still in in close proximity to the nuclear periphery. Co-staining experiments with α-Tubulin revealed that this cytoplasmic signal is closely associated with the microtubule manchette and it disappears from late elongating spermatids concomitant with manchette disassembly. In parallel, from the late round to early elongated stages onward, SUN5 accumulates at the HTCA region, where it ultimately persists in mature spermatozoa.

**Figure 6:**
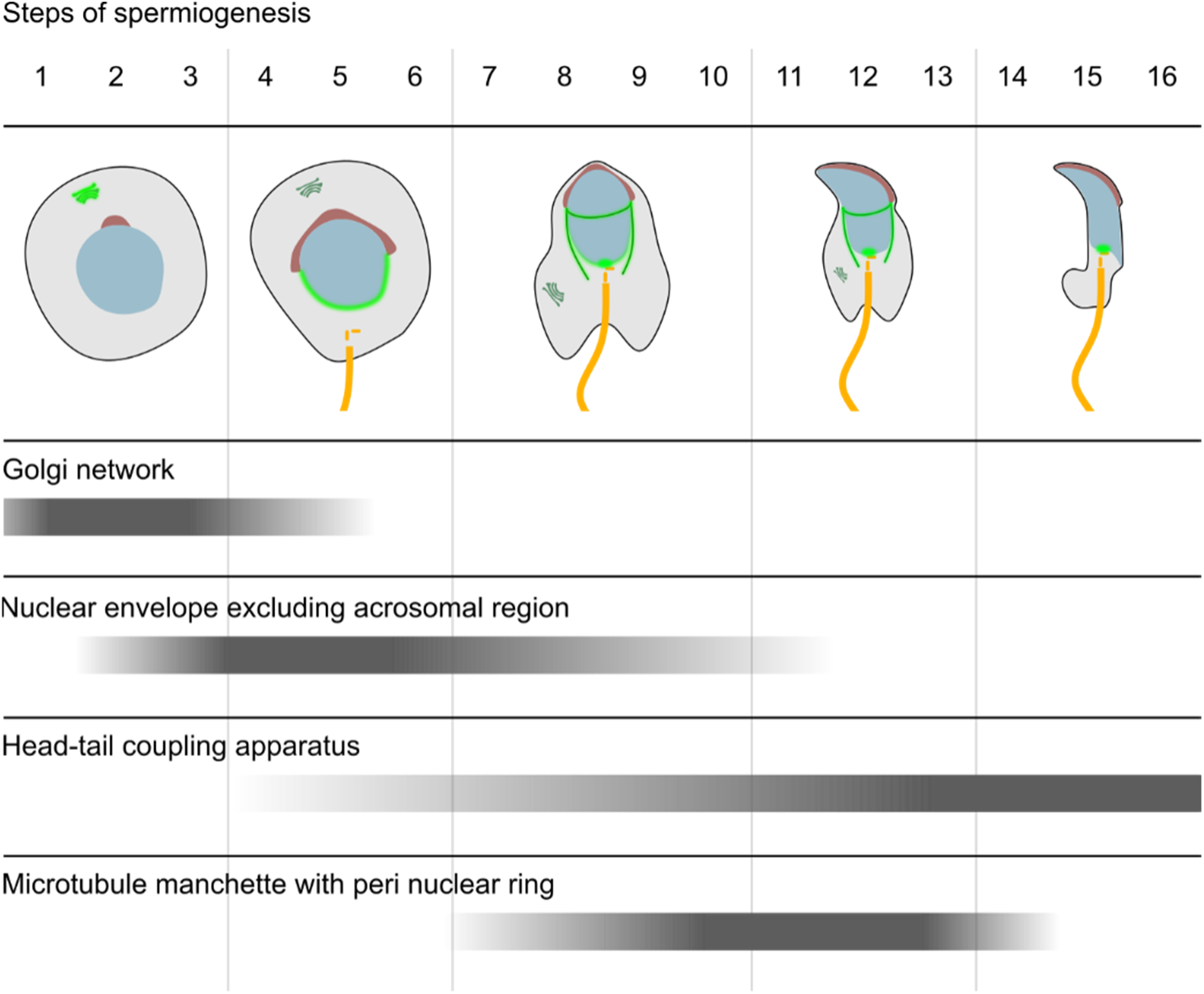
Model of SUN5 localization dynamics during spermiogenesis. Representative SUN5 localization patterns observed at different stages of spermiogenesis are summarized in this schematic model. The nucleus is depicted in blue, the acrosome in red, the sperm tail including the centriole pair in yellow and the Golgi apparatus in dark green. Light green indicates the dynamic SUN5 localization as observed in our U-ExM experiments. Grey bars represent the dynamic appearance and disappearance of SUN5 in relation to the depicted structure.

To analyze the SUN5 manchette association in more detail, we next performed U-ExM on SUN4 knock-out mice, taking advantage of the isolated manchettes that can be readily found in this model. Due to the absence of SUN4, spermatids from these mice lack a stable linkage between the manchette and the nucleus, resulting in dissociation of the manchette from the nuclear envelope. When not fully disassembled, the manchette persists as microtubule assemblies in the cytoplasm, spatially separated from the nucleus (Pasch et al., 2015; Calvi et al., 2015; Thoma et al., 2023).

As shown in Fig. 7, antibodies directed against the N-terminal domain and the PNS region of SUN5 both labeled the microtubule manchette, producing a predominantly punctate staining pattern indicative of multiple vesicular structures distributed along filaments. Co-staining with α-tubulin revealed that, in addition to the dotted signal along the manchette, a bright signal oriented perpendicular to the tubulin strands was observed at the anterior surface of the manchette. In contrast to the fluorescence along the microtubule filaments, the signals associated with the anterior surface exhibited a more continuous pattern. In the case of the antibodies targeting the PNS region, the anterior signal was markedly more intense than the staining of the core microtubule (MT) filaments (Fig 7b). Together with the analysis of wild-type spermatids (Fig. 5C, D; Fig. S2, Movie S1) these findings demonstrate that SUN5 is associated with the microtubule manchette and further suggests that the protein is present at the PNR.

**Figure 7:**
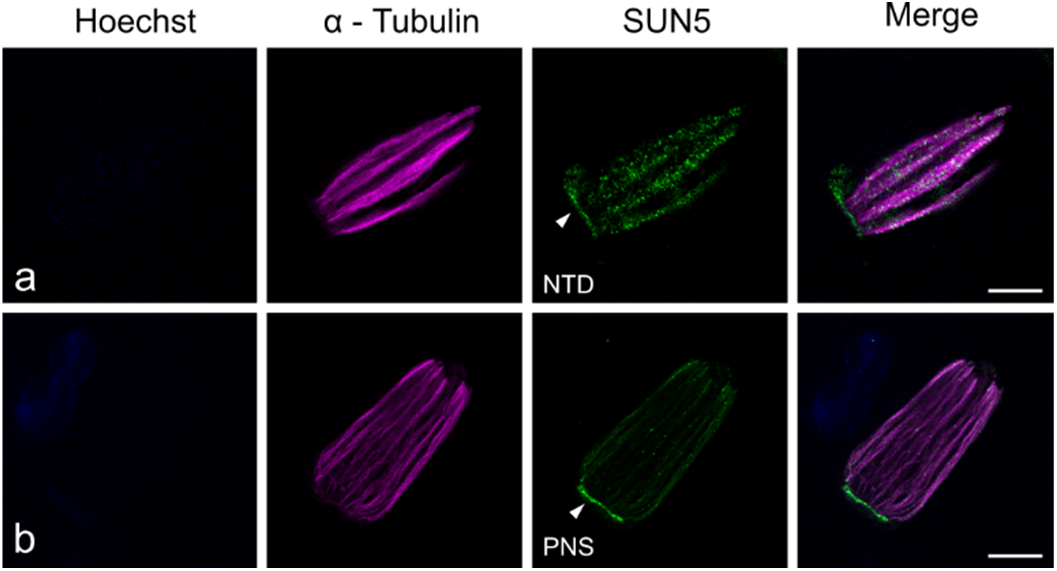
Association of SUN5 to the microtubule manchette and the PNR. Association of SUN5 to the microtubule manchette and the PNR. NE dissociated manchettes in spermatids from SUN4-ko mice were analyzed by U-ExM. Both, the anti-SUN5-NTD (a) and the anti-SUN5-PNS (b) clearly immunodecorated the manchette, which was costained with anti-α-tubulin antibodies. Slightly distant from the α-tubulin signal, the SUN5 antibodies aditionally recognized a perpendicular orientated structure, likely corresponding to the PNR. Scale bars: 10 µm; Images are representatives of at least four experimental repeats.

## Discussion

LINC complexes are traditionally described as molecular assemblies that physically connect the nucleoskeleton with the cytoskeleton (Crisp et al., 2006). In mammals, canonical LINC complexes are primarily formed by SUN1 and SUN2. They are broadly expressed in somatic cells and establish a characteristic arrangement in which SUN proteins anchored in the inner nuclear membrane (INM) interact with KASH partners in the outer nuclear membrane (ONM) (for comprehensive review see Starr and Fridolfsson, 2010). Interestingly, during spermiogenesis, the process by which haploid gametes differentiate into fertilization-competent spermatozoa (Kracklauer et al., 2013) three additional, testis-specific SUN proteins, SUN3, SUN4, and SUN5, are expressed, which appear to play central roles in this nuclear shaping and head-to-tail linkage (Göb et al., 2010; Calvi et al., 2015; Pasch et al., 2015; Zhu et al., 2016; Shang et al., 2017; Gao et al., 2020). SUN3 and SUN4 are part of the nuclear envelope-microtubule manchette junction, which serves as a crucial hub for sperm-specific nuclear shaping (Pasch et al., 2015; Calvi et al., 2015; Gao et al., 2020). Although initially presumed to be membrane proteins oriented to the cytoplasm (Shao et al., 1999), SUN4, and in consequence also SUN3, turned out to be, like SUN1 and SUN2, components of the inner nuclear membrane with their N-terminal domain facing the nucleoplasm and the C-terminal domains positioned within the PNS (Thoma et al., 2023).

### SUN5 shows a LINC-typical membrane topology in the HTCA region

In contrast to the other SUN proteins, relatively little was known about SUN5, the fifth member of the SUN-domain protein family in mammals. Moreover, the existing literature reported considerable inconsistencies concerning its localization and topology, which are crucial for interpreting the actual function of SUN5. While the authors of the original characterization of the protein reported a canonical localization (Frohnert et al., 2011), other, subsequent studies propose that the SUN domain of SUN5 is exposed to the cytoplasm (Zhang et al., 2021b; Zhang et al., 2024). Based on these and other findings in various model organisms, three distinct models of SUN5 topology and function have recently been discussed (Buglak et al., 2025). These include a classical, KASH dependent model, an NPC-related scenario and a direct model, which proposes that SUN5 binds directly to cytoplasmic interaction partners via a cytoplasm-facing SUN domain.

Based on our extensive analyses combining (a) detailed immunogold localization experiments using antibodies against epitopes presumed to reside on opposite membrane sides and (b) an *in vivo* protease protection assay, we now provide clear evidence that, within the HTCA region, SUN5 shares a membrane topology typical of SUN domain proteins, with its NTD facing the nucleoplasm and its CTD localized to the PNS. Thus, our findings strongly support a KASH-dependent, and potentially also an NPC-related, mechanism of HTCA linkage to the NE, while effectively excluding the model involving direct interaction of the SUN domain to cytosolic components. AlphaFold-based predictions of trimeric SUN5 assemblies estimate a structure length of approximately 12 nm from the transmembrane region to the end of the SUN domain. This fits well to our measurements of the INM–ONM spacing at the basal plate, which revealed a distance of 14.25 nm and is consistent with recently available data presented by Moecking et al. (2025, preprint), who measured a distance of 14.34 nm. Hence, the model in which SUN5 should span both nuclear membranes, as postulated by Zhang et al. (2024) and proposed as alternative model by Buglak et al. (2025) appears highly unlikely. This conclusion is further supported by the absence of membrane bulges in our electron micrographs, which would be expected if SUN5 were to bridge both nuclear membranes.

Taken together, our data combining direct immunogold labelling, ultrastructural distance measurements and AlphaFold predictions of SUN5 strongly support a classical LINC or NPC related model, in which SUN5 localizes to the inner nuclear membrane and presents its C-terminal domain within the PNS. These findings align well with the data presented in two recent preprints reporting cryo-electron microscopy analyses of the HTCA region in sperm of different mammals (Dendooven et al., 2025 preprint; Moecking et al., 2025 preprint). In both studies, regular protein lattices were observed at the HTCA, which are speculated to be formed by SUN5, thereby providing additional support for a classical LINC-dependent model. NPCs, however, did not appear to associate with these lattices, effectively ruling out the NPC-related model proposed by Buglak et al. (2025). This conclusion is further reinforced by previous analyses of NPC redistribution during spermiogenesis, which did not report any accumulation of NPCs at the HTCA (Ho, 2010).

In the past years, a number of potential interacting partners of SUN5 have been reported, among them Nxf1 and Nup 93 (He et al., 2024), DNAJB13 (Shang et al., 2018), CENTLEIN (Zhang et al., 2021b), Nesprin3 (Zhang et al., 2021a), LaminB1 and Septin12 (Zhang et al., 2024), most of which localize to or near the HTCA. A direct interaction between SUN5 and all of these candidates, however, would necessitate the presence of numerous concurrent binding sites. In addition, as discussed above, some of the proposed interactions are clearly inconsistent with the recent findings, particularly those suggesting a cytoplasmic interaction involving the SUN domain. As most studies describing such interactions employed GST-pulldown or co-IP assays to identify binding partners, it is possible that some of the reported associations are indirect rather then the result of direct physical interactions (De Las Rivas and Fontanillo, 2012). The currently prevailing model in literature proposes a mechanism in which SUN5 is localized to the NE at the HTCA, there connecting to PMFBP1 and SPATA6, which reside on the cytoplasmic side of the basal plate NE (Wu et al., 2020). However, no direct physical interaction between SUN5 and PMFBP1 has been demonstrated to date (Zhu et al., 2018), suggesting that the molecular composition of the head-to-tail linkage may be even more complex than currently assumed. Zhang and colleagues (2021b) identified CENTLEIN as the putative missing linker; however, it is also a cytoplasmic protein lacking a transmembrane domain. Our data clearly demonstrate that the SUN domain of SUN5 is not exposed to the cytosol, rendering such interactions and other similarly proposed models (Zhang et al., 2021b; Zhang et al., 2024; Buglak et al., 2025) very unlikely. Nevertheless, it is evident that CENTLEIN is critical for HTCA integrity (Zhang et al., 2021b) suggesting the involvement of another linker protein that spans the ONM and connects SUN5 to CENTLEIN. A promising candidate is Nesprin3, which has been reported to interact with SUN5 (Zhang et al., 2021a) and, as a member of the KASH protein family, would fit well with the idea of forming a canonical LINC complex. This hypothesis, however is challenged by the observation that Nesprin3 deficient mice are fertile (Ketema et al., 2013) and that Nesprin3 overtly interacts with SUN1. The subcellular localization and behavior of Nesprin3 appears to be linked to that of SUN1, which significantly differs to that of SUN5 during sperm differentiation (Göb et al., 2010; this study). Together, although some level of interaction may occur, it appears unlikely that Nesprin3 and SUN5 are functionally coupled. It should be noted here, that Lamin B1 has been identified as a nuclear binding partner of SUN5 (Zhang et al., 2024), but so far it remains the only known interaction partner in the nucleus. Beyond this, the available data in the literature on nuclear interactors are very limited. Taken together, while our findings refine the current model by excluding various cytoplasmic binding partners lacking a transmembrane domain, the precise interaction partners still need to be determined, and this issue should be systematically re-investigated in future studies.

### Dynamic SUN5 Behavior during sperm differentiation

Previous reports describing SUN5 at distinct subcellular locations suggest that its localization may be highly dynamic rather than static (Yassine et al., 2015). Supporting this notion, our detailed U-ExM analysis demonstrated a pronounced, stage-dependent redistribution of SUN5 during spermiogenesis, including its transient Golgi-association, its presence at the nuclear envelope, its accumulation at the HTCA as well as a previously unreported staining of the microtubule manchette and PNR (Fig. 6).

In early spermiogenesis, SUN5 shows partial association with GM130-positive structures, but with a spatial offset, suggesting a transient accumulation at the trans-Golgi rather than stable residence at the cis-Golgi network. This intermediate localization likely reflects trafficking of SUN5 towards the nuclear envelope and aligns with previous report describing SUN5 as a transient Golgi-associated protein in spermatids (Yassine et al., 2015). Subsequently, Golgi derived non-acrosomal vesicles bud from the Golgi network and fuse to the NE, leading to a gradual accumulation of SUN5 at the NE in early round spermatids. From this stage onward, the protein progressively concentrates into a single posterior spot at the HTCA. This signal persists in mature spermatozoa, marking the attachment site of the sperm tail and defining the final localization of SUN5, which is linked to its bona fide function in head-to-tail coupling (Shang et al., 2017; Zhang et al., 2021a; Zhang et al., 2024).

### Possible Functions of SUN5 Beyond Head-Tail Coupling

A rather surprising observation was that SUN5 redistribution appears to coincide with a progressive increase in cytoplasmic signal intensity during the transition from late round to early elongating spermatids. Using UxM, we could find that during this developmental stage a substantial fraction of SUN5 is closely associated with the microtubule manchette, with a tendency to accumulate at the PNR. This was further confirmed by the analysis of isolated manchettes in the SUN4 knockout model, where SUN5 was found to be enriched at microtubule assemblies that were detached from the nuclear envelope (Calvi et al., 2015; Pasch et al., 2015; Thoma et al., 2023). This observation suggests that SUN5 could function as a cargo of IMT and may there have additional functions, e.g. as a regulator of IMT.

IMT is a highly complex, spermiogenesis specific transport mechanism involved in delivering building blocksboth to the developing acrosome aswell asto posterior structureslike the HTCA and sperm tail. It shows many molecular parallels to intraflagellar transport (IFT), and its disruption can give rise to a wide range of pathological phenotypes, ultimately leading to male infertility. The microtubule manchette acts as the central hub of IMT, providing the structural basis both for the bi-directional transport mechanism and sperm head shaping and is physically coupled to the nucleus via the PNR (anterior) and SUN3/SUN4 containing LINC complexes (lateral) (Gao et al., 2025; Kierszenbaum, 2002; Kierszenbaum and Tres, 2004; Göb et al., 2010; Lehti and Sironen, 2016; Thoma et al., 2023; Miyata et al., 2024).

The pronounced staining of the manchette documented in our current study, both in intact cells and isolated manchettes, indicates that SUN5 might undergo IMT. This may represent a necessary transport route to its site of action, as direct access to the nuclear envelope becomes increasingly restricted during acrosome and manchette maturation, both progressively covering the nucleus. In a recent study examining the ultrastructural organization of the manchette, vesicles associated with the manchette surface were directly visualized (Judernatz et al., 2025), which fits well with our observation of punctate signals along the microtubule filaments of the manchette.

Beyond its association with the manchette, the pronounced signal detected at the PNR could reflect a more fundamental role in IMT or PNR function or stabilization. In support of this idea, Judernatz and colleagues (2025) reported that the PNR is closely associated with membranes in rat isolated manchettes, which may explain the presence of SUN5 as an integral membrane component in manchettes that are detached from the nuclei in SUN4-deficient spermatids. Importantly, this membrane-manchette association appears to be particularly robust, as it withstands even stringent extraction conditions. Further phenotypic evidence consistent with this concept comes from ultrastructural studies by Shang et al. (2017), who report disrupted biogenesis of the axoneme in SUN5 deficient mice. In this context, SUN5 might hold a critical role in shuttling vesicles from the manchette to the developing axoneme. However, this hypothesis requires direct experimental confirmation, representing an intriguing avenue for future research.

Overall, against this broader background, we here provide a comprehensive analysis of the subcellular localization and the dynamic behavior of SUN5, highlighting its stage-dependent redistribution and, for the first time, its specific association with the microtubule manchette and the PNR. Our findings offer new insights into the SUN5 network system and open up exciting directions for future studies on SUN5 functions beyond head-to-tail coupling.

## Acknowledgments

We thank Elisabeth Meyer-Natus and Dr. Fabian Link for introduction to the Tokuyasu method. We are grateful to Konstantin Wawra and Yeva Dzhokych for excellent technical assistance and thank Prof. Dr. Christian Stigloher and his team (Imaging Core Facility, Biocenter, University of Würzburg, Germany) for supporting the acquisition of EM data.

## Figure Legends

**Figure S1: Alpha-fold based model of a trimeric SUN5 assembly with local pLDDT values.**

**Figure S2: Cryo-section of an elongating spermatid probed with anti SUN4 and SUN5 antibodies.** SUN5 signal (green) appears on the anterior edge of the SUN4 (magenta) signal, indicating localization to the peri-nuclear ring (PNR), scalebar: 5 µm. Image is representative for at least four experimental repeats.

**Table S1: Primary Antibodies used in the presented study**

**Table S2: Secondary Antibodies used in the presented study**

**Code S1: FIJI-Macro to measure the width of the PNS**

**Code S2: FIJI-Macro to measure gold particle distance to the peri-nuclear space**

