## Supplementary Information for "Nanoscale imaging resolves canonical topology and intracellular dynamics of SUN5/SPAG4L during mammalian spermiogenesis"

**Supplementary information** to manuscript Herold et al.

**Table S1: Primary Antibodies used in the presented study**

| Antibody | Supplier | Catalog number | Dilution U-ExM / TEM |
|---|---|---|---|
| Mouse Anti-α-Tubulin | ABCD Antibodies (Geneva, Switzerland) | ABCD_AA344 | 1:50 |
| Mouse Anti-GM130 | BD Biosciences (Becton, Dickinson and Company, Franklin Lakes, NJ, USA) | 610822 | 1:100 |
| Rabbit Anti-SUN5-NTD | AG Alsheimer / SeqLab (SEQLAB Sequence Laboratories Göttingen GmbH, Göttingen, Germany) | --- | 1:100 / 1:400 |
| Guinea Pig Anti-SUN5-PNS | AG Alsheimer / SeqLab (SEQLAB Sequence Laboratories Göttingen GmbH, Göttingen, Germany) | --- | 1:100 / 1:400 |
| Rabbit Anti-SUN4-NTD | AG Alsheimer / SeqLab (SEQLAB Sequence Laboratories Göttingen GmbH, Göttingen, Germany) | --- | 1:100 |
| Guinea Pig Anti-SUN4-NTD | AG Alsheimer / SeqLab (SEQLAB Sequence Laboratories Göttingen GmbH, Göttingen, Germany) | --- | 1:100 |

**Table S2: Secondary Antibodies used in the presented study**

| Antibody | Supplier | Catalog number | Dilution |
|---|---|---|---|
| Alexa Fluor® 594 Donkey Anti-Guinea Pig | Jackson ImmunoReseatch larboatories Inc. (West Grove, PA, USA) | 706-585-148 | 1:100 |
| Alexa Fluor® 488 Donkey Anti-Rabbit | Jackson ImmunoReseatch larboatories Inc. (West Grove, PA, USA) | 711-545-152 | 1:100 |
| Alexa Fluor® 594 Donkey Anti-Rabbit | Jackson ImmunoReseatch larboatories Inc. (West Grove, PA, USA) | 711-585-152 | 1:50 |
| Alexa Fluor® 488 Goat Anti-Guinea Pig | Thermo Fischer Scientific (Waltham, MA, USA | A-11073 | 1:100 |
| Texas Red® Goat Anti-Mouse | Jackson ImmunoReseatch larboatories Inc. (West Grove, PA, USA) | 115-075-044 | 1:20 |
| 6nm gold Goat Anti-Rabbit | Dianova (Hamburg, Germany) | 111-195-144 | 1:10 |
| 12 nm gold Donkey Anti-Rabbit | Dianova (Hamburg, Germany) | 111-205-144 | 1:10 |
| 12 nm gold Donkey Anti-Guinea Pig | Jackson ImmunoReseatch larboatories Inc. (West Grove, PA, USA) | 706-205-148 | 1:10 |
| Peroxidase-conjugated Donkey Anti-Guinea Pig | Jackson ImmunoReseatch larboatories Inc. (West Grove, PA, USA) | 706-035-148 | 1:10000 |
| Peroxidase-conjugated Donkey Anti-Rabbit | Jackson ImmunoReseatch larboatories Inc. (West Grove, PA, USA) | 711-035-152 | 1:10000 |
| Peroxidase-conjugated Donkey Anti-Mouse | Jackson ImmunoReseatch larboatories Inc. (West Grove, PA, USA) | 115-035-146 | 1:10000 |

### Supplementary Images

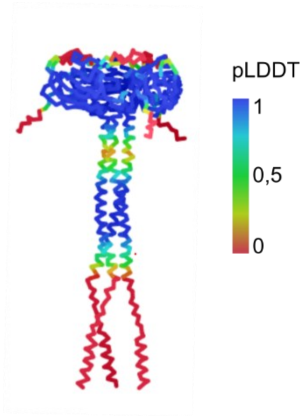


**Figure S1: Alpha-fold based model of a trimeric SUN5 assembly with local pLDDT values.**

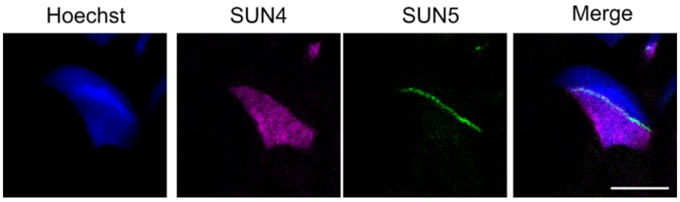


**Figure S2: Cryo-section of an elongating spermatid probed with anti SUN4 and SUN5 antibodies.** SUN5 signal (green) appears on the anterior edge of the SUN4 (magenta) signal, indicating localization to the peri-nuclear ring (PNR), scalebar: 5 µm. Image is representative for at least four experimental repeats.

### Code S1: FIJI-Macro to measure the width of the PNS

```
macro "PNS_Ausmessen_FWHM [q]" {

//settings for pre-processing (gauss), minimal allowed peak distance and FWHM
method (usually 0.5)

gaussSigma = 2;
minPeakDistance = 10;
fwhmThreshold = 0.5;
title = getTitle();

function smooth(array, sigma) {
  radius = round(2 * sigma);
  kernelSize = 2 * radius + 1;
  kernel = newArray(kernelSize);
  sum = 0;
  for (i = -radius; i <= radius; i++) {
    w = exp(-(i*i)/(2*sigma*sigma));
    kernel[i + radius] = w;
    sum += w;
  }
  for (i = 0; i < kernelSize; i++) kernel[i] /= sum;

  smoothed = newArray(lengthOf(array));
  for (i = 0; i < lengthOf(array); i++) {
    v = 0;
    for (j = -radius; j <= radius; j++) {
      k = i + j;
      if (k < 0) k = 0;
      if (k >= lengthOf(array)) k = lengthOf(array) - 1;
      v += array[k] * kernel[j + radius];
    }
    smoothed[i] = v;
  }
  return smoothed;
}

function findFWHM(profile, peakIndex, threshold) {
  peakVal = profile[peakIndex];
  half = peakVal * threshold;

  left = peakIndex;
  while (left > 0 && profile[left] > half)
    left--;

  right = peakIndex;
  while (right < lengthOf(profile)-1 && profile[right] > half)
    right++;

  return newArray(left, right);
}

// validate selection
getSelectionCoordinates(x, y);
if (x.length != 2) exit("Bitte eine gerade Linie mit dem Linien-Werkzeug
einzeichnen.");

profile = getProfile();
```

```
profile = smooth(profile, gaussSigma);

// normalize intensities
min = profile[0];
max = profile[0];
for (i = 1; i < lengthOf(profile); i++) {
  if (profile[i] < min) min = profile[i];
  if (profile[i] > max) max = profile[i];
}
normalized = newArray(profile.length);
for (i = 0; i < profile.length; i++)
  normalized[i] = (profile[i] - min) / (max - min);

// find peaks
peakIndices = newArray();
peakValues = newArray();
for (i = 1; i < lengthOf(normalized)-1; i++) {
  if (normalized[i] > normalized[i-1] && normalized[i] > normalized[i+1]) {
    peakIndices = Array.concat(peakIndices, i);
    peakValues = Array.concat(peakValues, normalized[i]);
  }
}
if (lengthOf(peakIndices) < 2) exit("Nicht genügend Peaks im Profil gefunden.");

max1Index = 0;
max2Index = 1;
if (peakValues[1] > peakValues[0]) {
  max1Index = 1;
  max2Index = 0;
}
for (i = 2; i < lengthOf(peakIndices); i++) {
  if (peakValues[i] > peakValues[max2Index]) {
    max2Index = i;
    if (peakValues[max2Index] > peakValues[max1Index]) {
      temp = max1Index;
      max1Index = max2Index;
      max2Index = temp;
    }
  }
}

peak1 = peakIndices[max1Index];
peak2 = peakIndices[max2Index];
peakToPeakDistance = abs(peak2 - peak1);
if (abs(peak2 - peak1) < minPeakDistance) exit("Gefundene Peaks zu nahe
beieinander.");
if (peak2 < peak1) {
  temp = peak1; peak1 = peak2; peak2 = temp;
}

fwhm1 = findFWHM(normalized, peak1, fwhmThreshold);
fwhm2 = findFWHM(normalized, peak2, fwhmThreshold);
left1 = fwhm1[0]; right1 = fwhm1[1];
left2 = fwhm2[0]; right2 = fwhm2[1];

// edgeMargin check
edgeMargin = 2; // pixel distance to edge

if (left1 <= edgeMargin || right1 >= lengthOf(normalized) - edgeMargin
    || left2 <= edgeMargin || right2 >= lengthOf(normalized) - edgeMargin) {
  exit("FWHM-Grenzen zu nah am Rand erkannt. Messung ungültig.");
```

```
}

pnsWidth = left2 - right1;
if (pnsWidth < 0) exit("Fehlerhafte Rand-Erkennung: PNS-Breite negativ.");
membrane1Width = right1 - left1;
membrane2Width = right2 - left2;
avgMembraneWidth = (membrane1Width + membrane2Width) / 2;

// display plot with markers and close the previous one
if (isOpen("Profil")) {
    selectWindow("Profil");
    run("Close");
}
Plot.create("Profil", "Position", "Normierte Intensität", normalized);
Plot.setColor("black");
Plot.setLineWidth(1);
Plot.setColor("red");
Plot.add("line", newArray(left1, left1), newArray(0, 1));
Plot.add("line", newArray(right1, right1), newArray(0, 1));
Plot.add("line", newArray(left2, left2), newArray(0, 1));
Plot.add("line", newArray(right2, right2), newArray(0, 1));
Plot.show();

// add values to the results table
row = nResults;
setResult("Bild", row, title);
setResult("Linie X1", row, x[0]);
setResult("Linie Y1", row, y[0]);
setResult("Linie X2", row, x[1]);
setResult("Linie Y2", row, y[1]);
setResult("PNS-Breite", row, pnsWidth);
setResult("Peak-Abstand", row, peakToPeakDistance);
setResult("Membrandicke 1", row, membrane1Width);
setResult("Membrandicke 2", row, membrane2Width);
setResult("Durchschnitt-Membrandicke", row, avgMembraneWidth);

updateResults();
}
```

**Code S2: FIJI-Macro to measure gold particle distance to the peri-nuclear space**

```
run("Clear Results");

// ===== Helper functions =====
function min(a, b) {
    return a < b ? a : b;
}
function max(a, b) {
    return a > b ? a : b;
}

// ===== Step 1: Sample the reference line ROI =====
// Look for ROI named "Membrane"
membraneIndex = -1;
nROIs = roiManager("count");
for (i = 0; i < nROIs; i++) {
    roiManager("Select", i);
    name = getInfo("roi.name");
    if (name == "Membrane") {
        membraneIndex = i;
        break;
    }
}

if (membraneIndex == -1) {
    exit("Membrane ROI not found!");
}

roiManager("Select", membraneIndex);
getSelectionCoordinates(lineX, lineY);

interpPerSegment = 100;
interpX = newArray();
interpY = newArray();

for (i = 0; i < lineX.length - 1; i++) {
    x1 = lineX[i];
    y1 = lineY[i];
    x2 = lineX[i+1];
    y2 = lineY[i+1];

    for (j = 0; j <= interpPerSegment; j++) {
        t = j / interpPerSegment;
        interpX[lengthOf(interpX)] = x1 + t * (x2 - x1);
        interpY[lengthOf(interpY)] = y1 + t * (y2 - y1);
    }
}

// ===== Step 2: Process ROIs by name and calculate euclidian distance  =====
goldSets = newArray("6nmI", "6nmO", "12nmI", "12nmO");
row = 0;

for (set = 0; set < goldSets.length; set++) {
    roiName = goldSets[set];
    roiFound = false;

    for (i = 0; i < nROIs; i++) {
        roiManager("Select", i);
        name = getInfo("roi.name");
```

```
            if (name == roiName) {
                roiFound = true;
                break;
            }
        }

        if (!roiFound) {
            print("Skipping " + roiName + " – ROI not found.");
            continue;
        }

        getSelectionCoordinates(particleX, particleY);
        nParticles = lengthOf(particleX);

        for (i = 0; i < nParticles; i++) {
            px = particleX[i];
            py = particleY[i];
            minDist = -1;

            for (j = 0; j < interpX.length; j++) {
                dx = px - interpX[j];
                dy = py - interpY[j];
                d = sqrt(dx*dx + dy*dy);
                if ((minDist < 0) || (d < minDist)) {
                    minDist = d;
                }
            }

            // Inside ROIs get negative distance
            if (roiName == "6nmI" || roiName == "12nmI") {
                minDist = -minDist;
            }

            // Save result
            setResult("Set", row, roiName);
            setResult("Point Index", row, i + 1);
            setResult("X", row, px);
            setResult("Y", row, py);
            setResult("Min Distance (px)", row, minDist);
            row++;
        }
    }

    run("Results...");
```
